# Integrated molecular analysis identifies a conserved pericyte gene signature in zebrafish

**DOI:** 10.1101/2021.09.13.459947

**Authors:** Yu-Huan Shih, Daneal Portman, Feston Idrizi, Ann Grosse, Nathan D. Lawson

## Abstract

Pericytes reside in capillary beds where they share a basement membrane with endothelial cells and regulate their function. However, little is known about embryonic pericyte development, in part, due to lack of specific molecular markers and genetic tools. Here, we applied single cell RNA-sequencing (scRNA-seq) of *platelet derived growth factor beta (pdgfrb)*-positive cells to molecularly characterize pericytes in zebrafish larvae. scRNA-seq revealed zebrafish cells expressing mouse pericyte gene orthologs while comparison to bulk RNA-seq from wild type and *pdgfrb* mutant larvae further refined a pericyte geneset. Subsequent integration with mouse pericyte scRNA-seq profiles revealed a core set of conserved pericyte genes. Using transgenic reporter lines, we validated pericyte expression of two genes identified in our analysis: *NDUFA4 mitochondrial complex associated like 2a* (*ndufa4l2a*), and *potassium voltage-gated channel, Isk-related family, member 4* (*kcne4*). Both reporter lines exhibited pericyte expression in multiple anatomical locations, while *kcne4* was also detected in a subset of vascular smooth muscle cells. Thus, our integrated molecular analysis revealed a molecular profile for zebrafish pericytes and allowed us to develop new tools to observe these cells *in vivo*.

**SUMMARY STATEMENT:** An integrated molecular analysis in zebrafish identifies new molecular markers for pericytes.

## INTRODUCTION

The vertebrate circulatory system comprises blood vessels and a heart, which are lined by a single layer of endothelial cells. The circulatory system delivers oxygen to tissues, while disposing of carbon dioxide and other waste products. Blood vessels also provide a conduit for immune cells and hormones of the endocrine system. Delivery and exchange of blood constituents to tissue is facilitated by capillary beds that are functionally tuned to their anatomical location (Holm et al., 2018). For example, in the central nervous system, capillaries of the blood brain barrier exhibit highly selective transport activity to limit entry of circulating factors that could damage neurons (Zhao et al., 2015). By contrast, liver and kidney capillary beds are important for filtering and removing serum components from circulation (Holm et al., 2018), requiring less restricted passage through the endothelial lining. In capillaries, vascular mural cells, known as pericytes, directly contact and share a basement membrane with endothelial cells and regulate their transport function (Holm et al., 2018). In the absence of pericytes, capillary function is altered. A notable example is the blood brain barrier where reduced pericyte coverage causes vascular leakage due to increased endothelial transcytosis (Armulik et al., 2010). Thus, pericytes serve as gatekeepers of capillary function within the circulatory system.

The pathways and mechanisms associated with pericyte-mediated organotypic capillary function have been well studied (Armulik et al., 2011; Holm et al., 2018). However, much less is known about embryonic pericyte development. Early studies in avian embryos revealed a dual origin for pericytes: most pericytes in the rostral regions of the central nervous system derived from neural crest, while elsewhere they arise from mesoderm (Etchevers et al., 2001; Korn et al., 2002). Subsequent studies in mouse and zebrafish suggest that this is a conserved aspect of pericyte development (Ando et al., 2019; Simon et al., 2012). A number of signaling pathways have also been implicated in early pericyte development, including Platelet derived growth factor beta receptor (Pdgfrb), which is highly expressed on pericytes, and its endothelial-expressed ligand (Pdgfb), along with components of the Notch signaling pathway (Ando et al., 2021; Dieguez-Hurtado et al., 2019; Hellstrom et al., 1999; Kofler et al., 2015; Levéen et al., 1994; Lindahl et al., 1997; Wang et al., 2014). However, little is known about the stepwise acquisition of pericyte identity during embryonic development. This knowledge gap stems, in part, from the lack of definitive molecular markers for pericytes in developing vertebrate embryos.

Numerous studies have focused on identifying pericyte-specific genes. Early efforts utilized microarray or RNA-seq performed on microvascular fragments from mutant backgrounds that reduce the numbers of pericytes or incorporated transgenic lines to isolate vascular mural cell populations (Bondjers et al., 2006; Bondjers et al., 2003; He et al., 2016; Jung et al., 2018). More recently, application of scRNA-seq has begun to reveal new markers for pericytes (Vanlandewijck et al., 2018). However, these efforts have largely focused on adult mouse tissues and studies at embryonic stages, or in other vertebrate models, are lacking. In the zebrafish embryo, which has proven useful for investigating pericyte development, only a few pericyte-specific genes have been identified, limiting the utility of this model (Ando et al., 2016; Ando et al., 2021; Ando et al., 2019). To address this issue, we have applied scRNA-seq on *pdgfrb*-positive cells isolated from zebrafish larvae, along with parallel integration with related datasets, to better define a pericyte gene signature. We subsequently leveraged our findings to develop new reporter lines to directly visualize pericytes *in vivo*.

## RESULTS AND DISCUSSION

Transgenic zebrafish lines made using the *pdgfrb* locus drive fluorescent protein expression in pericytes and numerous non-vascular cell types (Ando et al., 2016; Vanhollebeke et al., 2015), making definitive identification of pericyte markers challenging with bulk RNA-seq. Therefore, to identify pericyte-expressed genes, we performed 3’ single-cell RNA-sequencing (scRNA-seq) using Egfp-positive cells isolated by fluorescent activated cell sorting (FACS) from 5 dpf larvae bearing *TgBAC(pdgfrb:egfp)*^*ncv22*^ (Ando et al., 2016). After filtering and integrated analysis of replicate libraries, we identified more than 50 distinct clusters from 12,865 cells (**Fig. S1A, B**), underscoring the heterogeneity of *pdgfrb:egfp*-positive cells. We next used selected cluster-specific genes to assign cell identities (**Fig. S1C, Table S1.1, 1.2; see Methods**). For example, *actinodin1* (*and1*) defined epidermal cells, while *mitfa* identified melanocytes and neurons expressed *elavl3* (**Fig. S1C, Table S1.1, 1.2**; (Kim et al., 1996; Lawson et al., 2020; Lister et al., 1999; Zhang et al., 2010)). We could not assign identities to ten clusters, although cells in these cases exhibited high expression of collagens (*col1a2* and *col12a1a*), suggesting fibroblast or mesenchymal characteristics (Rajan et al., 2020). Surprisingly, most clusters displayed very low levels of endogenous *pdgfrb* transcript (**Fig. S1C, Table S1.3**), suggesting that these cells may have been isolated due to ectopic *pdgfrb:egfp* expression. Alternatively, *pdgfrb:egfp* expression in progenitor cells and Egfp perdurance may have contributed to their capture by FACS. Given high *pdgfrb:egfp* levels in pericytes (Ando et al., 2016), we focused on clusters expressing higher relative levels of endogenous *pdgfrb* (**Fig. 1A**, *pdgfrb*^*hi*^ cells, log_2_ average expression > 0.6; **Table S1.3**) and smooth muscle cell clusters, based on their gene expression profiles being similar to pericytes (Vanlandewijck et al., 2018).

**Figure 1.**
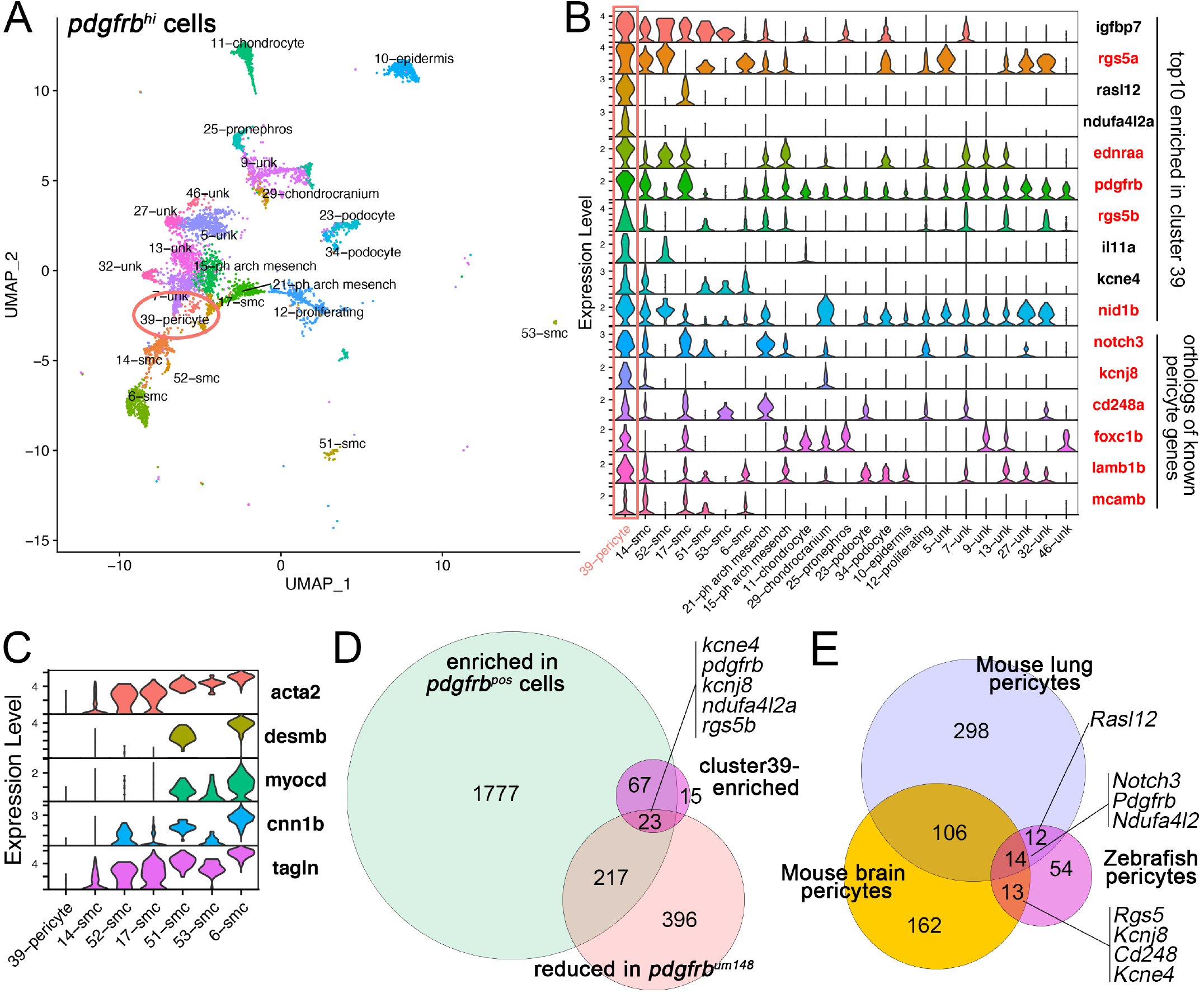
scRNA-seq analysis of *pdgfrb:egfp*-positive cells. **(A)** UMAP plot of *pdgfrb:egfp*-positive cells isolated from *TgBAC(pdgfrb:egfp)*^*ncv34*^ larvae at 5 dpf. Only clusters with the highest *pdgfrb* levels are shown. Pericyte cluster is circled. **(B)** Violin plot of top 10 enriched genes from cluster 39 (sorted by decreasing log_2_ fold change relative to all other clusters, adjp<0.05) and other selected pericyte genes (log_2_ fold change>0.75, adjp<0.05). Known pericyte genes in red. Pericyte cluster is highlighted by a rectangle. **(C)** Violin plot showing known smooth muscle genes expressed in indicated clusters. **(D)** Venn diagram of genes enriched in *pdgfrb:citrine*-positive cells (compared to negative cells, log_2_ fold change>1, adjp<0.05), reduced in *pdgfrb:citrine*-positive *pdgfrb*^*um148*^ mutant cells (compared to wild type; log_2_ fold change<-1, adjp<0.05), and enriched in cluster 39 (log_2_ fold change>0.75, adjp<0.05). **(E)** Venn diagram of genes enriched in mouse brain and lung pericytes and zebrafish pericytes. Only genes with log_2_ fold change>0.75, adjp<0.05 in each respective individual analysis were considered.

Among *pdgfrb*^*hi*^ cells, cluster 39 expressed the highest *pdgfrb* levels (**Table S1.3**). Moreover, mouse orthologs for half of the ten most enriched genes in this cluster (ranked by log_2_ fold change, adj.p<0.05; **Table S1.4**) are expressed in pericytes, including duplicates of *regulator of G protein signaling 5* (*rgs5a* and *rgs5b*), *endothelin receptor type Aa* (*ednraa*), and *nidogen 1b* (*nid1b*; **Fig. 1B, Table S1.4**; (Bondjers et al., 2003; Kitazawa et al., 2011; Sakhneny et al., 2021)). Other significantly enriched genes (log_2_ fold change>0.75, adjp<0.05) included *notch3*, which is expressed in zebrafish brain pericytes (Wang et al., 2014) and *potassium inwardly rectifying channel subfamily J member 8* (*kcnj8*), a marker of mouse brain pericytes ((Bondjers et al., 2006), **Fig. 1B, Table S1.4**). We similarly noted cluster-enriched expression of *endosialin/CD248 molecule a* (*cd248a*), *forkhead box c1b* (*foxc1b*), *melanoma cell adhesion molecule b* (*mcamb*), and *laminin b1b* (*lamb1b*), orthologs of which are expressed in mouse pericytes (Bagley et al., 2008; Middleton et al., 2005; Sakhneny et al., 2021; Siegenthaler et al., 2013), **Fig. 1B, Table S1.4**). Consistent with pericyte identity, cells in cluster 39 did not express *smooth muscle actin alpha 2* (*acta2*), a definitive marker of zebrafish smooth muscle cells (SMCs; (Georgijevic et al., 2007; Whitesell et al., 2014)), or other SMC genes such as *desminb* (*desmb*), *myocardin* (*myocd*), *calponin1b* (*cnn1b*) or *transgelin* (*tagln*, **Fig. 1C**; (Georgijevic et al., 2007; Wang et al., 2014; Whitesell et al., 2019)). Thus, cluster 39 likely comprises pericytes and we hereafter refer to genes enriched in these cells as “pericyte genes”.

To further characterize the expression of pericyte genes, we incorporated bulk RNA-seq analysis of *TgBAC(pdgfrb:citrine)*^*s1010*^ cells (Lawson et al., 2020). We considered two differential gene sets where pericyte-specific genes would be expected to be found. First, genes enriched in wild type *pdgfrb:citrine*-positive compared to - negative cells (log_2_ fold change>1, adjp<0.05; **Table S2.1**). Second, genes reduced in *pdgfrb:citrine*-positive cells from *pdgfrb*^*um148*^ mutants, which lack pericytes (Ando et al., 2021), compared to wild type siblings at 5 dpf (log_2_ fold change<-1, adjp<0.05; **Table S2.2**). Intersection of these datasets revealed 23 out of 105 pericyte genes with enrichment in *pdgfrb:citrine* cells and reduction in *pdgfrb*^*um148*^ mutants (**Fig. 1D, Table S2.3, S3**), including *interleukin 11a* (*il11a*), *NDUFA4 mitochondrial complex associated like 2a* (*ndufa4l2a*), and *potassium voltage-gated channel, Isk-related family, member 4* (*kcne4*), which have not been previously characterized pericyte-expressed. Most pericyte genes (90 out of 105), including many previously identified orthologs, exhibited enrichment in *pdgfrb:citrine*-positive compared to -negative cells, but were not reduced in *pdgfrb*^*um148*^ mutants (**Fig. 1D, Table S3**). This is likely due to their widespread expression in other *pdgfrb*-positive cell types not affected in *pdgfrb*^*um148*^ mutant larvae. We would note in this regard that pericytes make up less than 1% of all *pdgfrb*-expressing cells (**Fig. S1A; Table S1.2**). *Pdgfrb* itself is likely detected as downregulated despite its broad expression in non-pericyte cell types since the *um148* allele causes non-sense mediated decay of the mutant transcript (Kok et al., 2015).

An additional consideration we made for zebrafish pericyte genes was whether they were conserved among vertebrate models. Therefore, we incorporated scRNA-seq analysis of mouse pericytes (He et al., 2018; Vanlandewijck et al., 2018). Re-clustering datasets from brain and lung vascular cells identified pericyte populations characterized by high levels of *Cspg4, Pdgfrb, Notch3, Mcam* and *Ifitm1* and low expression of SMC markers (*Myocd* and *Cnn1*; **Fig. S2A-D, Table S4**). Intersection of mouse and zebrafish pericyte genes revealed a large degree of overlap (39/93 zebrafish genes found in mouse pericytes; intersection based on mouse orthologs, see **Material and Methods**; **Fig. 1E, Table S3**). Notably, the degree of overall overlap between zebrafish and mouse pericytes from either tissue was comparable to that between mouse pericytes from brain and lung. This analysis identified a core set of 14 pericyte genes that were commonly enriched in pericytes across the three datasets, including *Notch3* and *Pdgfrb* (**Fig. 1E, Table S3**). From this core gene set, all displayed significant enrichment in *pdgfrb*^*pos*^ cells by bulk RNA-seq, but only 2 (*pdgfrb* and *ndufa4l2a*) showed reduction in *pdgfrb*^*um148*^ mutants (**Table S3**), again suggesting most of these genes were not restricted to pericytes. This is consistent with a high proportion of non-pericyte cells expressing many of the pericyte genes from our scRNA-seq analysis (*pct*.*2* value in **Table S1.4**). Nonetheless, these observations demonstrate that zebrafish larval pericytes share a conserved molecular signature with those in adult mouse tissues.

Together, our molecular analysis establishes a gene signature for zebrafish pericytes at the larval stage. To demonstrate the utility of this dataset for studying zebrafish pericytes, we chose two previously uncharacterized candidate pericyte genes for generation of transgenic reporter lines. The first was *ndufa4l2a*, which is among the most consistently enriched and specific genes from our analysis (**Table S3**). To assess the expression pattern of *ndufa4l2a*, we constructed a recombinant bacterial artificial chromosome (BAC) with super folder green fluorescent protein (sfGFP) inserted into the first exon and used this to generate a stable transgenic line (*TgBAC(ndufa4l2a:sfgfp)*^*um382*^). Confocal imaging of the brain vasculature in *TgBAC(ndufa4l2a:sfgfp)*^*um382*^ larvae at 5 dpf revealed expression in cells closely associated with blood vessels throughout the mid- and hindbrain (**Fig. 2A**). Cells expressing *ndufa4l2a:sfgfp* also co-expressed *abcc9:gal4ff;uas:rfp* (referred to hereafter as *abcc9:rfp*), a known pericyte marker (Ando et al., 2019; Vanlandewijck et al., 2018), and appeared to wrap around brain blood vessels, consistent with pericyte morphology (**Fig. 2A, B**). We also observed expression of *ndufa4l2a:sfgfp* in *abcc9:rfp*-positive pericytes lining retinal endothelial cells (**Fig. 2C**). In the trunk, *ndufa4l2a:sfgfp*-positive cells expressed *abcc9:rfp* and were closely associated with intersegmental vessels at 5 dpf (**Fig. 2D**). We did not detect sfGFP expression in vascular mural cells along the dorsal aorta (**Fig. 2D**), most of which express *acta2* at this stage consistent with their identity as vascular SMCs (VSMCs; (Whitesell et al., 2014)). In both the trunk and cranial vasculature, we noted *ndufa4l2a:sfgfp*-positive cells that did not express *abcc9:rfp* (arrowheads denoted by asterisks in **Fig. 2A, D**). This is likely due to mosaic somatic silencing that frequently occurs when using the Gal4/UAS system (Goll et al., 2009). Thus, the *ndufa4l2a:sfgfp* appeared largely restricted to *abcc9*-positive pericytes throughout the vasculature. However, we also noted non-vascular expression throughout the epidermis, as well as cells with a mesenchymal appearance on the surface of the lower jaw (**Fig. S3A, B**). Interestingly, none of the epidermis clusters from scRNA-seq analysis expressed *ndufa4l2a* (**Fig. S1C**), suggesting the observed *ndufa4l2a:sfgfp* expression is ectopic.

**Figure 2.**
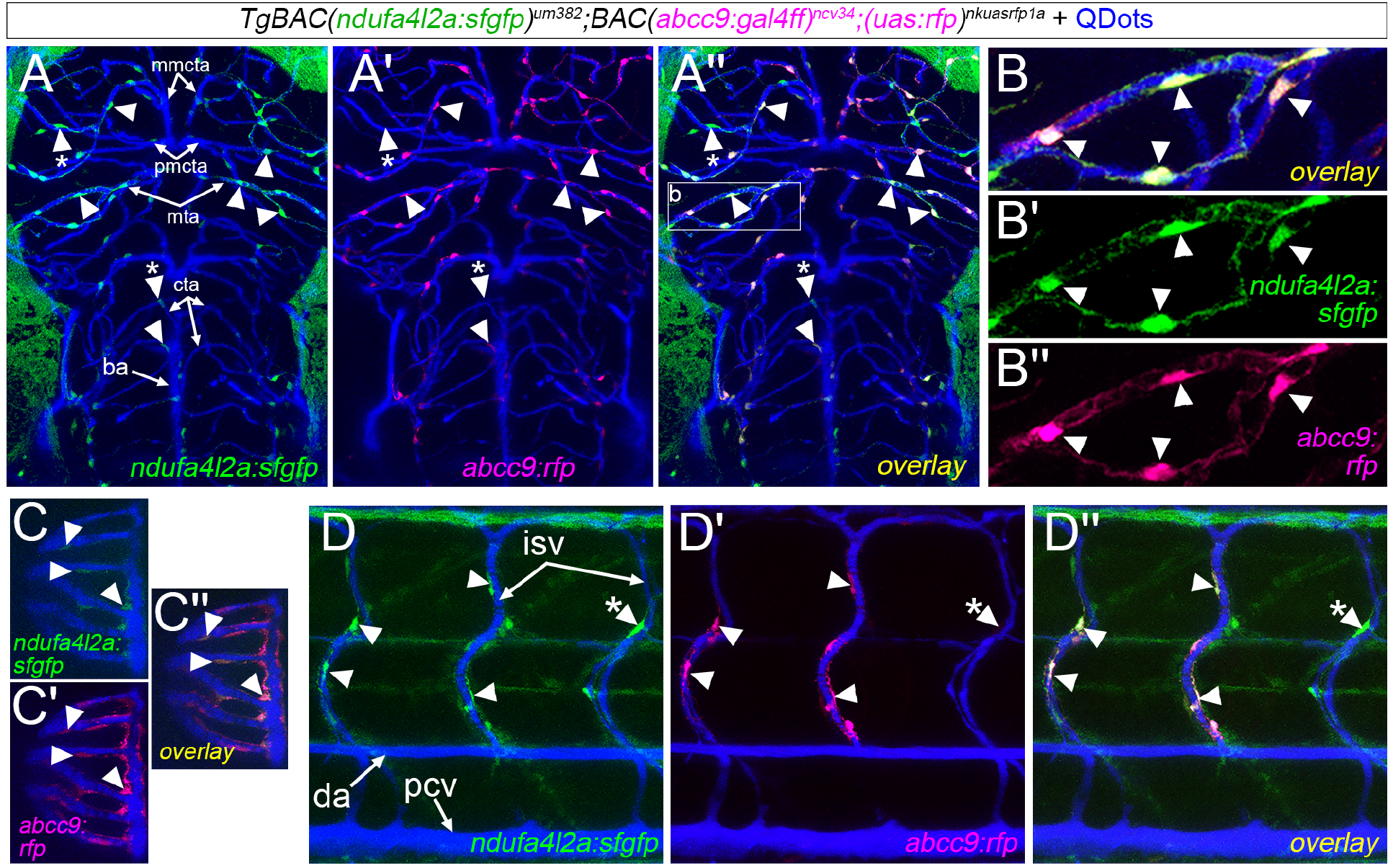
The *ndufa4l2a* locus drives pericyte-specific expression. **(A-D)** Vertical projections of confocal stacks of *TgBAC(ndufa4l2a:sfgfp)*^*um382*^;*(abcc9:gal4ff)*^*ncv34*^;*(uas:rfp)*^*nkuasrfp1a*^ larvae at 5 dpf subjected to angiography with QDots. **(A)** Expression of *ndufa4l2a:sfgfp* in pericytes (selected cells denoted by arrowheads) along branches emanating from the middle mesencephalic central artery (mmcta), and posterior mesencephalic central artery (pmcta), and along the metencephalic artery (mta). Positive cells are also seen along central arteries (cta). **(A’)** *abcc9:rfp* expression and **(A’’)** overlay for same embryo as **(A)**; box indicates magnified area in **(B)**. Asterisks denote cells only expressing sfGFP. **(B-B’’)** Magnified image of *ndufa4l2a:sfgfp* and *abcc9:rfp* co-expression in pericytes (denoted by arrows). **(C-C’’)** Expression of **(C)** *ndufa4l2a:sfgfp* and **(C’)** *abcc9:rfp* in retinal pericytes. **(C’’)** Overlay; arrowheads denote zebrafish co-expressing pericytes. **(D)** Expression of **(D)** *ndufa4l2a:sfgfp* and **(D’)** *abcc9:rfp* in pericytes along the intersegmental vessels (isv). **(D’’)** Overlay; arrows denote co-expressing pericytes; asterisk indicates cells only expressing sfGFP. da is dorsal aorta, pcv is posterior cardinal vein. **(A-C)** Dorsal views, anterior is up. **(D)** Lateral view, dorsal is up, anterior is to the left.

In addition to *ndufa4l2a*, we also generated a reporter line using a BAC encompassing the *kcne4* locus (*TgBAC(kcne4:sfgfp)*^*um333*^). Similar to the *ndufa4l2a:sfgfp* transgene, *kcne4:sfgfp* expression was present in brain pericytes, in this case co-expressed with a *pdgfrb:gal4ff* transgene driving *uas:rfp* (**Fig. 3A**). We also observed *kcne4:sfgfp* expression in *pdgfrb*-positive pericytes in the retinal vasculature (**Fig. 3B**) and along the ISVs and dorsal aorta in the trunk (**Fig. S3C**). Unlike endogenous *ndufa4l2a*, which was restricted to pericytes in *pdgfrb*-positive cells by scRNA-seq analysis, *kcne4* was also enriched in SMCs (**Fig. 1B; Table S1.1**). However, we did not observe *kcne4:sfgfp* expression in *acta2:mcherry*-positive VSMCs in association with the dorsal aorta in the trunk or the Circle of Willis in the brain (**Fig. 3C, D**). In both cases *kcne4:sfgfp* could be detected in pericytes in association with nearby blood vessels (**Fig. 3C, D**). In the intestine, which is surrounded by SMCs, we observed scattered *kcne4:sfgfp*-positive cells that co-expressed *acta2:mcherry*, along with *acta2*-negative cells in direct contact with blood vessels consistent with pericyte morphology (**Fig. 3E**). In contrast to the dorsal aorta, *kcne4:sfgfp* was expressed in *acta2*-positive VSMCs in aortic arch vessels branching from the ventral aorta (VA) and in the cardiac ventricle, but not along the ventral aorta (VA) itself (**Fig. 3F, G**). sfGFP-positive cells that appear on the ventral aorta (VA) in Fig. 3F and G are located ventrally and are not directly associated with blood vessels (**Fig. S3D, E**). We also detected *acta2*-negative cells with pericyte morphology that expressed *kcne4:sfgfp* along the hypobranchial artery, although these cells did not appear prominently in other vessels in this region (**Fig. 3F**). Thus, *kcne4:sfgfp* is expressed in pericytes, as well as anatomically restricted subsets of SMCs in the gut and VSMCs in the head. As with the *ndufa4l2a:sfgfp* transgene, we also detected epidermal expression in *TgBAC(kcne4:sfgfp)*^*um333*^ larvae (**Fig. S3C**). Taken together, these transgenic lines underscore the utility of our molecular analysis and provide new genetic tools for studying pericytes in zebrafish.

**Figure 3.**
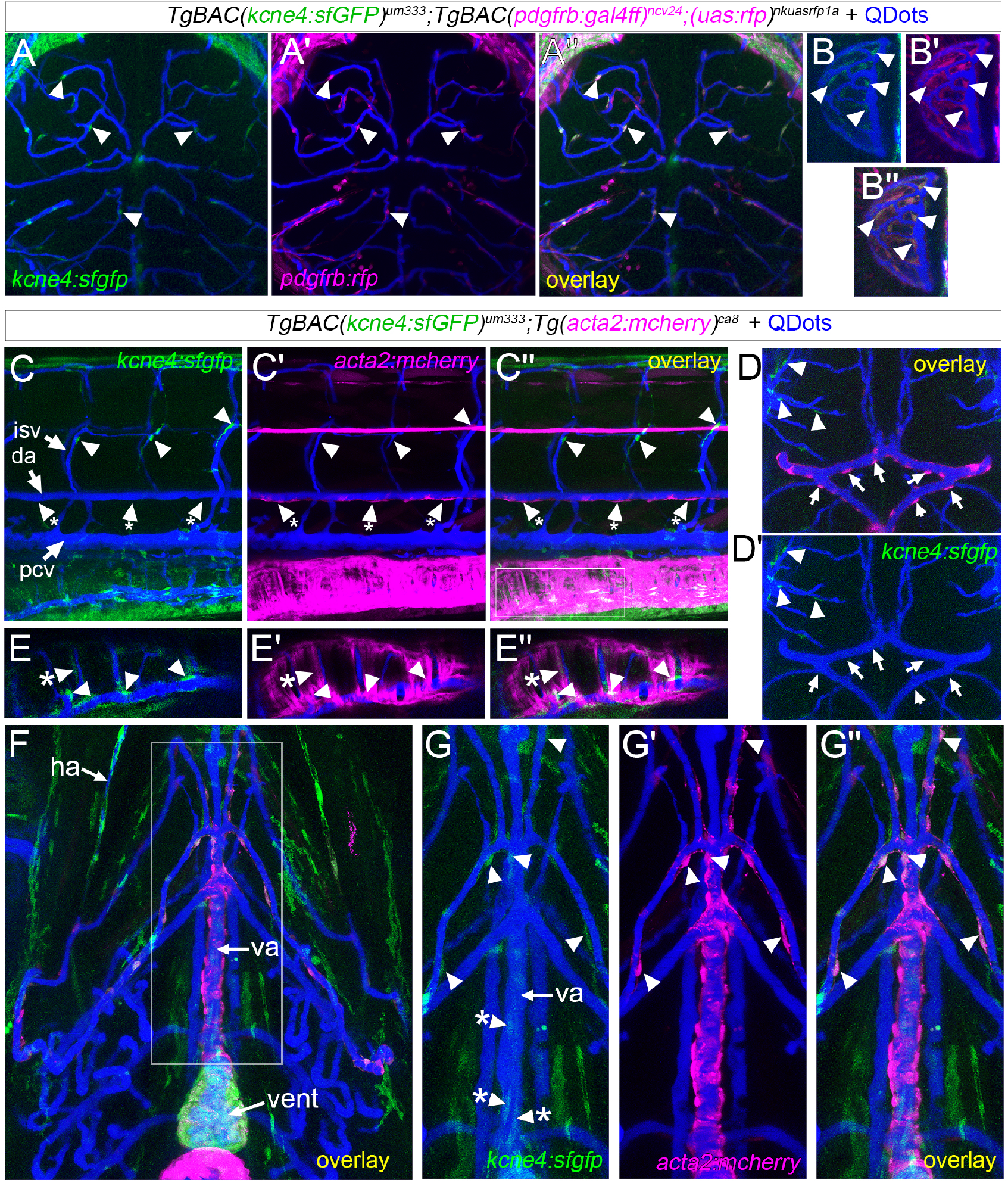
The *kcne4* locus drives expression in pericytes and selected vascular smooth muscle cells. **(A-G)** Vertical projections of confocal stacks of *TgBAC(kcne4:sfgfp)*^*um333*^ larvae at 5 dpf subjected to angiography with QDots also bearing **(A**,**B)** *TgBAC(pdgfrb:gal4ff)*^*ncv24*^;*(uas:rfp)*^*nkuasrfp1a*^ (referred to as *pdgfrb:rfp*) or **(C-G)** *Tg(acta2:mcherry)*^*ca8*^. **(A)** Expression of *kcne4:sfgfp* in pericytes (selected cells denoted by arrowheads) in cranial vessels. **(A’)** *pdgfrb:rfp* expression and **(A’’)** overlay for same embryo as **(A). (B-B’’)** Expression of **(B)** *kcne4:sfgfp* and **(B’)** *pdgrb:rfp* in retinal pericytes. **(B’’)** Overlay; arrowheads denote co-expressing pericytes. **(C-C’’)** Expression of **(C)** *kcne4:sfgfp* in pericytes (arrows) along the intersegmental vessels (isv) and **(C’)** *acta2:mcherry* in vascular smooth muscle cells (VSMCs; arrowheads with asterisk) on the dorsal aorta (da). **(C’’)** Overlay, pcv is posterior cardinal vein. **(D-D’)** Imaging of the Circle of Willis. **(D)** Overlay image showing *acta2:mcherry*-positive VSMCs (arrows) and nearby cranial arteries with *kcne4:sfgfp*-positive pericytes (arrowheads). **(D’)** *kcne4:sfgfp* channel only, showing absence of expression in VSMCs. **(E-E’’)** Single confocal section of the intestine; magnified from area denoted in **(C’’). (E)** *kcne4:sfgfp* expression is detected in pericytes on the intestinal vessels (arrowheads) and **(E-E’)** scattered *acta2:mcherry*-positive smooth muscle cells (arrowhead denoted by asterisk). **(E’’)** Overlay of **(E)** and **(E’). (F-G)** Ventral view of outflow tract and associated vasculature. **(F)** Overlay image showing *kcne4:sfgfp* expressed in pericytes on the hypobranchial artery (ha). Ventral aorta (va) is noted, as is the cardiac ventricle (vent). **(G-G”)** Area of magnification shown by box in (F). Channels and overlay as indicated in each panel. Plain arrowheads denote *kcne4:sfgfp* expression in *acta2*-positive VSMCs. Arrowheads marked with an asterisk are sfGFP-positive cells located ventral to the ventral aorta (va; see **Fig. S3D, E**). **(A-B, D)** Dorsal views, anterior is up. **(C, E)** Lateral view, dorsal is up, anterior is to the left. **(F-G)** Ventral views, anterior is up.

The zebrafish has long been a valuable model for studying vascular development, providing numerous novel insights into conserved cellular and molecular processes that contribute to blood vessel growth and function. However, most of these efforts have focused on endothelial cells, facilitated by a wide range of technical resources for their study in the zebrafish embryo, including numerous transgenic tools, mutants, and a broad catalog of molecular markers. By contrast, only a handful of studies have addressed the development of zebrafish pericytes and genetic tools in this regard are limited. Our current study begins to fill these gaps by establishing a molecular signature comprised of more than 100 genes expressed in zebrafish larval pericytes. We demonstrate the utility of this dataset by establishing reporter lines for visualizing pericytes in zebrafish larvae. Together, our findings will provide a foundation for identifying functional candidate genes and further developing genetic tools to dissect the genetic requirements for pericyte development during embryogenesis.

## MATERIALS AND METHODS

### Zebrafish Lines and Maintenance

All studies were performed under the auspices of animal protocols approved by the University of Massachusetts Medical School Institutional Animal Care and Use Committee. *TgBAC(pdgfrb:citrine)*^*s1010*^, *TgBAC(pdgfrb:egfp)*^*ncv22*^, *pdgfrb*^*um148*^, *TgBAC(pdgfrb:gal4ff)*^*ncv24*^, *TgBAC(abcc9:gal4ff)*^*ncv34*^, *Tg(uas:rfp)*^*nkuasrfp1a*^, *Tg(acta2:mcherry)*^*ca8*^ have been described elsewhere. Lines generated in this study are described below.

### RNA-seq

For bulk RNA-seq, *citrine*-positive cells from *TgBAC(pdgfrb:citrine)*^*s1010*^;*pdgfrb*^*um148*^ mutant larvae at 5 dpf were obtained in the course of previous studies in parallel to cells from wild type larvae (Lawson et al., 2020; Whitesell et al., 2014). Mutant siblings were identified and separated based on the absence of brain pericytes by confocal microscopy. Larval dissociation, FACS isolation (UMass Medical School Flow Cytometry Core), RNA isolation, and library construction were applied as described previously (Lawson et al., 2020; Quillien et al., 2017; Whitesell et al., 2014). For single-cell RNA-seq, wild type *TgBAC(pdgfrb:egfp)*^*ncv22*^ larvae at 5 dpf were dissociated as described previously (Lawson et al., 2020; Quillien et al., 2017; Whitesell et al., 2014), except cells were not fixed. Live cells were subjected to FACS to isolate EGFP-positive cells and approximately 10,000 cells were loaded onto a 10x Chromium (10x Genomics) for generation of single cell droplets. We constructed 3’ 10x scRNA-seq libraries according to the manufacturer’s recommendations (v3, 10x Genomics) and we performed pilot depth sequencing on a MiSeq (Illumina; UMass Medical School Deep Sequencing Core Labs) to verify quality and captured cell numbers. Subsequently, libraries were sequenced to greater depth on a HiSeq2500 (Illumina; Genewiz).

### Computational Analysis

All pipelines were run remotely at the Massachusetts Green High Performance Computing Center (MGHPCC). We performed mapping of bulk RNA-seq using a DolphinNext pipeline as described previously (Lawson et al., 2020). Reads were mapped onto GRCz11 and expression levels quantified using our custom transcript annotation (V4.3.2, (Lawson et al., 2020)). We identified differentially expressed genes using DeSeq2 run in DEbrowser, as previously (Lawson et al., 2020). Reads from scRNA-seq libraries were mapped onto GRCz11 using Cell Ranger (v3.1.0, 10x Genomics) in a DolphinNext environment and gene quantifications were made using the V4.3.2 annotation as previously (Lawson et al., 2020). We utilized Seurat (v4; (Hao et al., 2021)) for clustering using the output from Cell Ranger. We used Rstudio in the Open OnDemand environment at MGHPCC to run Seurat as previously described (Lawson et al., 2020). We applied integrated analysis to combine two replicate scRNA-seq libraries from *pdgfrb:egfp* cells (R script for *pdgfrb:egfp* analysis available in supplementary **File S1**). Cells were filtered for feature counts and mitochondrial proportion (see **File S1**), yielding a total of 12,865 cells across the two libraries. Following integration and identification of common anchors, Uniform Manifold Approximation and Projection (UMAP) was applied to reduce dimensionality based on the first 40 principal components and clusters were identified using a resolution 2. We identified all cluster-specific markers and used these to assess cluster identity by manual comparison to whole mount *in situ* hybridization patterns available through (ZFIN) www.zfin.org. To identify putative pericyte cell clusters, we performed text-based searching of all cluster-specific genes for selected genes previously identified as pericyte-expressed. To integrate bulk RNA-seq with the pericyte gene sets from scRNA-seq analysis, we used the top 105 pericyte-enriched genes (log_2_FC>0.75 relative to all other clusters, adj.p<0.05), along with those enriched in *pdgfrb:citrine*-positive cells versus -negative (log_2_FC>1, adj.p<0.05) and reduced in *pdgfrb:citrine pdgfb*^*um148*^ cells compared to wild type (log_2_FC<-1, adj.p<0.05). To obtain mouse pericyte gene sets, we used published data downloaded from GEO as raw count tables (GSE99235, GSE98816 (He et al., 2018; Vanlandewijck et al., 2018)). Each dataset was individually analyzed using Seurat run as above (see **File S2** for commands and parameters for clustering). Pericyte clusters were identified based on expression of marker genes identified previously (Vanlandewijck et al., 2018). Pericyte gene sets were generated from the top enriched genes (log_2_ fold change>0.75 relative to all other clusters, adj.p<0.05). To identify mouse orthologs for zebrafish pericyte genes, we utilized two data sources: a curated HomoloGene dataset from the Mouse Genome Informatics database (http://www.informatics.jax.org/downloads/reports/HOM_AllOrganism.rpt) and zebrafish:mouse orthology from ZFIN (https://zfin.org/downloads/mouse_orthos.txt). From 105 zebrafish pericyte genes, we identified 93 with mouse orthologs. Two mouse orthologs, *Lamb1* and *Rbpms2*, are orthologous to pericyte genes that are duplicated in zebrafish. Gene sets were intersected to generate Venn diagrams based on gene symbol using intervene (Khan and Mathelier, 2017) with the following commands: --type list --save-overlaps. Proportional Venn diagrams were generated with resulting data using eulerr (https://github.com/jolars/eulerr).

### BAC construction and transgenesis

Bacterial artificial chromosome (BAC) plasmids containing *kcne4* (DKEY-16C17) or *ndufa4l2a* (DKEY-11F4) loci were purchased from Source BioScience and verified for presence of target sequence by PCR for the first coding exon (exon 2 for *kcne4*, exon 1 for *ndufa4l2a*; primers kcne4-e1-F and kcne4-e1-R, or ndufa4l2a-e1-F and ndufa4l2a-e1-R, see **Table S5** for primer sequences). Recombinant BACs were constructed based on established protocols using pRedET (GeneBridges, Bussmann, 2011 #145}). An Isce-Tol2-Tn5neo cassette was amplified from pGEM-Isce-Tol2-Tn5-neo (Addgene #176619) by PCR using pBelo_pGEMT-F and pBelo_pGEMT-R (**Table S5**), gel purified and electroporated into bacteria carrying the target BAC and pRedET. Correctly targeted clones were identified by PCR. We next introduced the sfGFP coding sequence with an optimized Kozak consensus sequence into the first coding exon of each locus. A template plasmid (p-sfGFP-FRT-amp-FRT, Addgene #176620) was constructed by inserting a gBlock encoding sfGFP followed by an SV40 polyA sequence and a FRT-flanked ampicillin cassette in place of EGFP in the pEGFP-C1 plasmid (Clontech). For *kcne4* and *ndufa4l2a* BACs, we amplified the sfGFP/amp cassette using primers kcne4_sfGFP_f and kcne4_FRT-Amp_R, or ndufa4l2a_sfGFP_F1 and ndufa4l2a_FRT-Amp_R1 (**Table S5**), respectively. Templates were gel-purified and electroporated into bacteria bearing the target BAC and pRedET. Following recombination, correct clones were identified by PCR and the ampicillin resistance cassette was removed using the 707-FLPe plasmid according to recommended instructions (GeneBridges). Recombinant BACs were purified using PureLink™ HiPure Plasmid Midiprep Kit (ThermoFisher Scientific) and injected into 1-cell stage zebrafish embryos together with I-SceI (NEB). Only embryos exhibiting low mosaic sfGFP expression were grown to adulthood. We identified a single founder for each recombinant BAC. These lines are referred to as: *TgBAC(kcne4:sfgfp)*^*um333*^ and *TgBAC(ndufa4l2a:sfgfp)*^*um382*^. Imaging of larval stages was performed by confocal microscopy as described elsewhere (Ando et al., 2021).

## ACKNOWLEDGEMENTS

We thank members of the Lawson Lab for helpful comments on the manuscript. We thank John Polli and Patrick White for their efforts in fish care and facility maintenance. We thank Naoki Mochizuki and Koji Ando for providing transgenic lines used in this study.

## COMPETING INTERESTS

The authors declare no competing interests.

## FUNDING

These studies were supported by R21NS105654 (NINDS) and R35HL140017 (NHLBI) from the National Institutes of Health awarded to N. D. L.

## DATA AVAILABILITY

Raw and processed files for *pdgfrb:egfp* scRNAseq and bulk RNA-seq of *pdgfrb*^*um148*^ mutants are available at GEO (GSE176129). Bulk RNA-seq datasets from wild type *pdgfrb:citrine*-positive and -negative cells were previously published and are available in GEO (GSE152759), as is mouse scRNA-seq (GSE99235 and GSE98816).

## SUPPLEMENTARY FILES

**File S1**. R script with Seurat commands for analysis of zebrafish *pdgfrb:egfp* scRNA-seq.

**File S2**. R script with Seurat commands for analysis of mouse pericyte scRNA-seq. Includes notes on different parameters used for brain and lung datasets.

**Table S1**. Cluster-enriched genes from *pdgfrb:egfp* scRNA-seq analysis.

**Table S2**. DeSeq2 analysis of *pdgfrb:citrine* bulk RNA-seq datasets.

**Table S3**. Integrated bulk and scRNA-seq output for zebrafish pericyte genes.

**Table S4**. Output from FindAllMarkers (Seurat) for mouse scRNA-seq analysis.

**Table S5**. Oligonucleotide primers used in this study.

## Notes

### Competing Interest Statement

The authors have declared no competing interest.

