## Supplementary material for "Integrated molecular analysis identifies a conserved pericyte gene signature in zebrafish": File S2

# commands for clustering mouse vascular cells
# NCBI GEO accession numbers:
# GSE98816 brain vascular cells; GSE99235 lung vascular cells
# use raw read counts for each
# following is for lung vascular cells; identical commands used for brain, except for 
# number of PCs used for clustering

MmLungCellsRawCounts <- read.table("GSE99235_Lung_samples_raw_reads_counts_matrix.txt", 
                                    fill = TRUE, header = TRUE, row.names = 1, sep = "", as.is = TRUE)

MmLungCells = CreateSeuratObject(counts = MmLungCellsRawCounts, project = "MmLungVascSc")
MmLungCells <- NormalizeData(MmLungCells, normalization.method = "LogNormalize", scale.factor = 10000)
MmLungCells <- FindVariableFeatures(MmLungCells, selection.method = "vst", nfeatures = 2000)

#scale data
all.genes <- rownames(MmLungCells)
MmLungCells <- ScaleData(MmLungCells, features = all.genes)
MmLungCells = RunPCA(MmLungCells, features = VariableFeatures(object = MmLungCells), npcs = 50)

# ID optimal PCs
# use same approach for brain cells
# see https://hbctraining.github.io/scRNA-seq/lessons/sc_exercises_clustering_analysis.html
# Determine percent of variation associated with each PC
pct <- MmLungCells[["pca"]]@stdev / sum(MmLungCells[["pca"]]@stdev) * 100
# Calculate cumulative percents for each PC
cumu <- cumsum(pct)
# Determine which PC exhibits cumulative percent greater than 90% and % variation associated with the PC as less than 5
co1 <- which(cumu > 90 & pct < 5)[1]
co1
# PC where contribution is only 5% to standard deviation
#[1] 43
# next step determines where percent change in variation between consecutive PCs is less than 0.1%
# Determine the difference between variation of PC and subsequent PC
co2 <- sort(which((pct[1:length(pct) - 1] - pct[2:length(pct)]) > 0.1), decreasing = T)[1] + 1
# last point where change of % of variation is more than 0.1%.
co2
# PC where percent change in variation between consecutive PCs is less than 0.1%
# [1] 13
# Minimum of the two calculation
#pcs <- min(co1, co2)
#pcs
#[1] 13
# for brain vascular cells PC=16

MmLungCells <- FindNeighbors(MmLungCells, dims = 1:13)
MmLungCells <- FindClusters(MmLungCells, resolution = 0.5)
MmLungCells <- RunUMAP(MmLungCells, dims = 1:13)

DimPlot(MmLungCells, reduction = "umap", label = TRUE)+NoLegend()

#markers to define clusters for reordering
DefaultAssay(MmLungCells) = "RNA"
VlnPlot(MmLungCells, 
        features = c("Cspg4","Pdgfrb","Notch3","Mcam","Ifitm1","Myocd","Cnn1","Cdh5","Lum","Dcn"), 
        stack = TRUE, flip=TRUE, combine = FALSE) + NoLegend()

########################################################
# reorder and rename clusters
#0,4,8 - endothelial
#1,2,3,7 - fibroblast
#5-pericyte
#6 - VSMC

MmLungCellsForClusterReID = MmLungCells 
#change ordering first
newClstOrder = c(5,1,7,3,2,6,8,0,4)
Idents(MmLungCellsForClusterReID) = factor(Idents(MmLungCellsForClusterReID), levels=newClstOrder)

#rename clusters with cell types and original cluster numbers
clusterNames = c("pericyte","1-fibroblast","7-fibroblast","3-fibroblast",
                 "2-fibroblast","vsmc","8-endothelial","0-endothelial","4-endothelial")
names(clusterNames) = levels(MmLungCellsForClusterReID)
MmLungCellsForClusterReID = RenameIdents(MmLungCellsForClusterReID, clusterNames)

DimPlot(MmLungCellsForClusterReID, reduction = "umap", label = TRUE, repel = TRUE)+NoLegend()

#markers to define clusters/for Supp Figure
DefaultAssay(MmLungCellsForClusterReID) = "RNA"
VlnPlot(MmLungCellsForClusterReID, 
        features = c("Cspg4","Pdgfrb","Notch3","Mcam","Ifitm1","Myocd","Cnn1","Cdh5","Lum","Dcn"), 
        stack = TRUE, flip=TRUE, combine = FALSE) + NoLegend()

#top pericyte genes from zebrafish, exclusive of Notch3 and Pdgfrb
VlnPlot(MmLungCellsForClusterReID, 
        features = c("Igfbp7","Rgs5","Rasl12","Ndufa4l2","Ednra",
                     "Il11","Kcne4","Nid1","Pcdh18"), 
        stack = TRUE, flip=TRUE, combine = FALSE) + NoLegend()

MmLungMarkers = FindAllMarkers(MmLungCellsForClusterReID, only.pos = TRUE, min.pct = 0.25, logfc.threshold = 0.5)
write.csv(MmLungMarkers, "LungAllMarkersRes05.csv")
