## Supplementary Figures for "Integrated molecular analysis identifies a conserved pericyte gene signature in zebrafish"

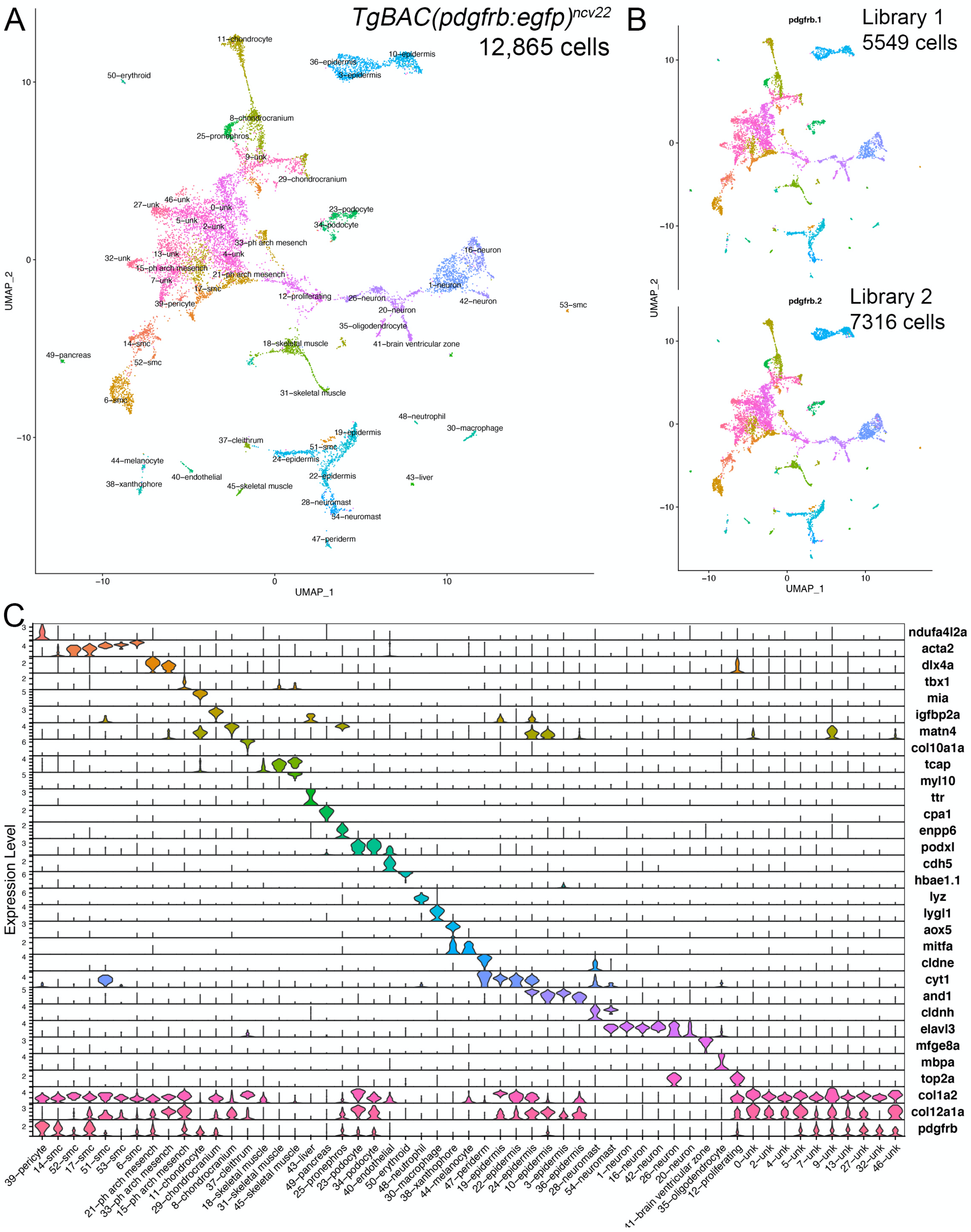


**Figure S1. Initial characterization of scRNA-seq from *pdgfrb:egfp*-positive cells. (A)** UMAP plot of *pdgfrb:egfp*-positive cells isolated from *TgBAC(pdgfrb:egfp)$^{ncv34}$* larvae at 5 dpf. Only clusters passing appropriate filters are shown (see **File S1**). Names of cell clusters are indicated. smc – smooth muscle cell; unk – unknown; ph arch mesench – pharyngeal arch mesenchyme. **(B)** UMAP plot of same cells as in **(A)** but split based on replicate libraries. **(C)** Stacked violin plot of selected cluster-enriched genes ($\log_2$ fold change>0.75, adjp<0.05) used to assign cell identity.

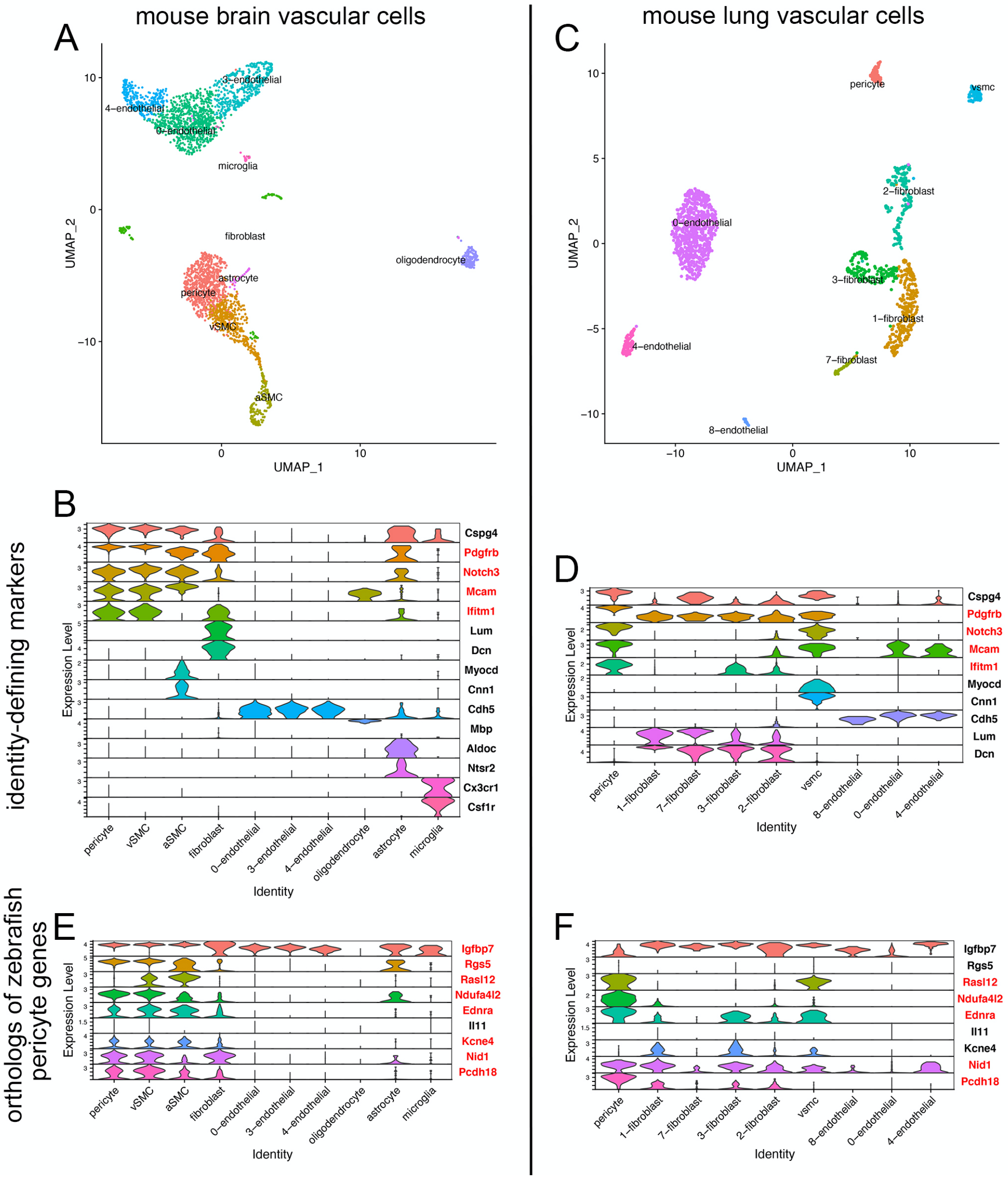


**Figure S2. scRNA-seq analysis of mouse pericytes from brain and lung. (A, C)** UMAP plots of vascular cells from **(A)** brain or **(C)** lung. See **File S2** for clustering parameters. **(B, D)** Stacked violin plot of selected cluster-enriched genes ($\log_2$ fold change>0.75, adjp<0.05) used to assign cell identity for vascular cells from **(B)** brain or **(D)** lung. **(E, F)** Stacked violin plot showing expression of mouse orthologs for top 10 enriched genes from pericytes in zebrafish. Red denotes significant enrichment ($\log_2$ fold change>0.75, adjp<0.05) in pericytes from **(E)** brain or **(F)** lung pericytes.

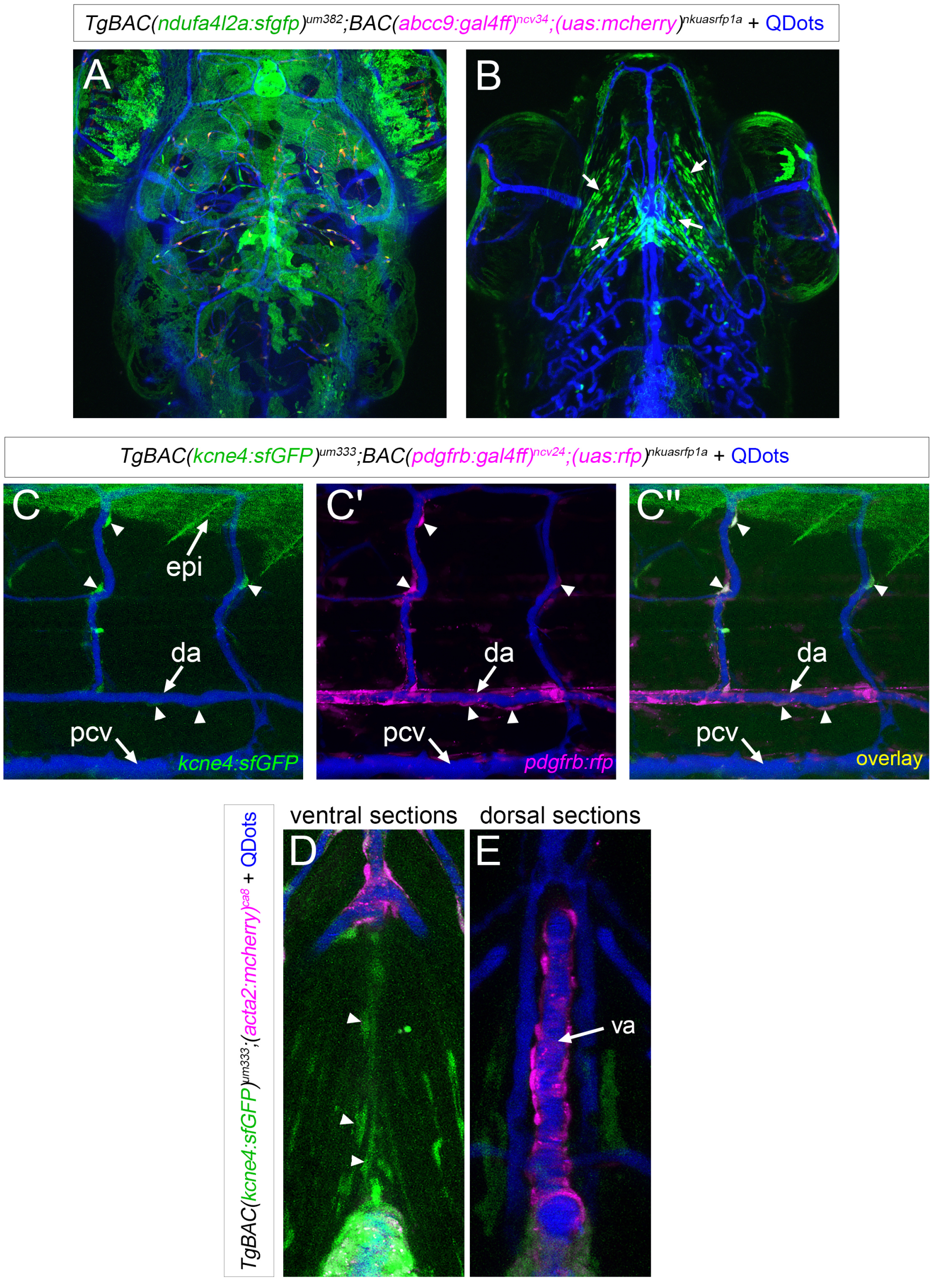


**Figure S3. Characterization of *TgBAC(kcne4:sfGFP)*$^{um333}$ and *TgBAC(ndufa4l2a:sfGFP)*$^{um382}$ larvae**. **(A-E)** Vertical projections of confocal stacks from larvae at 5 dpf subjected to angiography with QDots (blue). **(A, B)** *TgBAC(ndufa4l2a:sfGFP)*$^{um382}$;*BAC(pdgfrb:gal4ff)*$^{ncv24}$*;(uas:rfp)*$^{nkuasrfp1a}$ larvae; **(A)** Dorsal view showing extensive sfGFP expression on the epidermal surface of the head. **(B)** Ventral view showing sfGFP-positive cells with mesenchymal appearance in the lower jaw. These cells do not appear to be directly associated with blood vessels. **(C)** *TgBAC(kcne4:sfGFP)*$^{um333}$; *TgBAC(pdgfrb:gal4ff)*$^{ncv24}$*;(uas:rfp)*$^{nkuasrfp1a}$ larva, lateral view, anterior to the left, dorsal is up. Arrowheads denote kcne4:sfGFP-positive cells co-expressing RFP. da – dorsal aorta, pcv – posterior cardinal vein, epi – epidermis, isv – intersegmental vessel. **(D, E)** Ventral views of a *TgBAC(kcne4:sfGFP)*$^{um333}$*;(acta2:mcherry)*$^{ca8}$ larva. Each set of vertical projections was constructed from a single stack of confocal sections taken of the same larva. **(D)** Ventralmost sections, including epidermis to just below the ventral aorta, which is not included in this projection. Arrowheads denote mesenchymal *kcne4:sfGFP*-positive cells located ventral to the ventral aorta **(E)** Vertical projection of sections just dorsal to those in **(D)** showing ventral aorta (va) covered with *acta2:mcherry*-positive vascular smooth cells.
